# Sample-based regulatory intervention for managing risk within heterogeneous populations, with phytosanitary inspection of mixed consignments as a case study

**DOI:** 10.1101/441543

**Authors:** Stephen E. Lane, Rob M. Cannon, Anthony D. Arthur, Andrew P. Robinson

## Abstract

Inspection of consignments of imported goods is commonly undertaken at national borders in order to prevent incursions of pests and diseases and deter malefactors. Inspection of the whole consignment is usually either impossible or inefficient, so inspection of a random sample is used instead.

The size of the random sample for plant products is justified by appeal to International Standards for Phytosanitary Measures No. 31, “Methodologies for Sampling of Consignments”. ISPM 31 notes that “A lot to be sampled should be a number of units of a single commodity identifiable by its homogeneity […]” and “Treating multiple commodities as a single lot for convenience may mean that statistical inferences can not be drawn from the results of the sampling.”

However, commonly consignments are heterogeneous, either because the same commodities have multiple sources or because there are several different commodities. The ISPM 31 prescription creates a substantial impost on border inspection because it suggests that heterogeneous populations must be split into homogeneous sub-populations from which separate samples of nominal size must be taken.

We demonstrate that if consignments with known heterogeneity are treated as stratified populations and the random sample of units is allocated proportionally based on the number of units in each stratum, then the nominal sensitivity at the consignment level is achieved if our concern is the level of contamination in the entire consignment taken as a whole. We argue that unknown heterogeneity is no impediment to appropriate statistical inference. We conclude that the international standard is unnecessarily restrictive.

## 1 Introduction

### 1.1 Background

Border biosecurity programs are integral to the protection of our biodiverse natural environments, social amenity, and the economy through prevention of the entry of invasive pests and diseases. The economic cost (either directly, or from control measures) of invasive species has been estimated to be AUD 13.6 billion in Australia (Hoffmann and Broadhurst, 2016), up to NZD 3.3 billion in New Zealand (Giera and Bell, 2009), CND 34.5 billion in Canada (Colautti et al., 2006) and over USD 200 billion in the United States (Pimentel, 2011).

Border inspection for biosecurity is typically the responsibility of National Governments and is carried out for verifying the effectiveness of pre-arrival treatments, the detection of material that may pose a biosecurity risk, to gather information about contamination rates, and to deter any potential wrongdoing. Such pre-border and border intervention on a range of imported goods is based on the risk profile of the goods and international agreements.

It is often impractical to inspect all items in a consignment, so only a sample is inspected. In general a consignment would be deemed compliant only if no contaminated units are found in the sample, and non-compliant otherwise. For examples of sampling in the regulatory context, see Robinson (2017) and Venette et al. (2002).

The number required to be sampled is set to provide a certain probability (the sensitivity) that at least one contaminated item would be detected from the sample, given a particular prevalence of contaminated items, or less often, given a specified number of contaminated items. The Binomial distribution can be used for large consignments to determine this number. As an example, a typical application sets a prevalence (referred to as a design prevalence) at 0.5% and calculates the sample size required to have a 95% chance (the sensitivity) of detecting at least one contaminated item. In this case the required sample is 598, which is universally rounded to 600 for convenience. Ideally the design prevalence and sensitivity are chosen to provide an acceptable level of residual risk. When the regulator applies this approach, they are accepting that for consignments that do have a prevalence of infested items of 0.5%, in 5% of consignments no contaminated items will be found and these consignments will pass inspection. This example will be used throughout this paper to provide a tangible example of some concepts.

Formally, denote the design prevalence by *p*, the desired sensitivity by *S_d_*, and the number of units to be inspected by *n*. The regulator sets the parameters *p* and *S_d_*, then determines the number of units to be sampled (*n*), so that the probability that one or more contaminated units is found is greater than *S_d_*. For large consignments we can use the Binomial distribution to obtain the sensitivity 
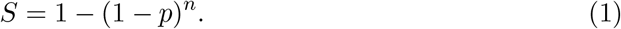

Expressing Equation (1) in terms of *n* gives us the (minimum) number of units to sample to achieve the desired sensitivity *S_d_*, as: 
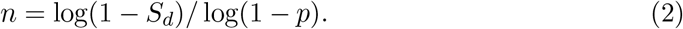

Historically, this sampling occurs within single lines in a consignment; a line comprises a single commodity. Consignments may however include multiple lines, either different commodities or the same commodity from different growers. It is natural to assume that identical commodities from different growers might have different levels of contamination. This expectation, combined with the misapprehension that a simple random sample of a consignment with likely heterogeneity would not achieve the desired level of sensitivity, appears to have resulted in the following recommendation under ISPM 31 (International Plant Protection Convention, 2008) on the topic of heterogeneous consignments (lots) of plant products:

“A lot to be sampled should be a number of units of a single commodity identifiable by its homogeneity in factors such as: origin, grower, packing facility, species, variety, degree of maturity, exporter, area of production, regulated pests and their characteristics, treatment at origin, or type of processing.

The criteria used by the NPPO to distinguish lots should be consistently applied for similar consignments.

Treating multiple commodities as a single lot for convenience may mean that statistical inferences can not be drawn from the results of the sampling.”

This prescription implies that in order for a heterogeneous consignment to satisfy the regulatory biosecurity requirements based on achieving a desired level of sensitivity (e.g. 95%) and a given design prevalence (e.g. 0.5%), it must be split into its homogeneous lines, and these must each be subjected to, for example, the 600 unit sample.

In what follows we consider that the contamination rate of the consignment as a whole is equal to the design prevalence, accepting that the rate within different parts of the consignment might be higher or lower than this value, and show that if the sample is split proportionately between the different parts, the sensitivity is at least as high as the value derived based on a single homogeneous consignment.

### 1.2 This Paper

The goal of this paper is to demonstrate that ISPM 31’s recommendation against mixing heterogeneous lines (lots) is unnecessarily restrictive, and that there are ways of sampling mixed lines that do achieve the required sensitivity against contamination without increasing the number of units we need to include in the sample.

Some critical assumptions are still required. First, we assume that the regulator is happy to apply their compliance rule to the entire consignment, in other words the entire consignment will only be deemed compliant if the sample taken from the consignment returns no contaminated items. Under this assumption the regulator is not specifically worried about higher levels of contamination in some lines, as long as the overall contamination rate of the consignment satisfies their design target. However, under this approach, if contamination is detected in any of the units sampled, then all of the lines from the consignment must be rejected. Second, our solution involves treating the lines in the consignment as if they were strata. We assume that once the sample is split, the required number of units from each line are randomly selected from the respective lines. We show that the act of stratifying the consignment by line and then allocating the total inspection sample (e.g. the 600 unit sample) proportionally to the stratum population counts will deliver nominal sensitivity (at least 95%) against a given overall contamination rate (0.5% as an example). Jointly, these arguments suggest that ISPM 31 is too restrictive in its prescription for mixed consignments.

## 2 ISPM 31 and Heterogeneity

The sole statistical reference provided for the ISPM 31 sample size calculations is Cochran’s 1977 Sampling Techniques (Cochran, 1977), and the calculations themselves can be located within a body of work called “design-based sampling theory”. Importantly, there is no statistical constraint or requirement for homogeneity of a sampled population within design-based sampling theory (Cochran, 1977). Indeed, samples are commonly collected and analysed across substantially heterogeneous populations, such as human and economic populations in official statistics, and forest communities in natural resource management. The only constraints are (i) that the sample be taken according to one of a number of different kinds of random sample designs, for example as detailed in ISPM 31, and (ii) if contamination is detected in any of the units sampled, then *all* of the lines from which samples were taken must be rejected. If the heterogeneity is unknown within a single diverse line then a simple random sample will deliver nominal sensitivity by design.

## 2.1 Dividing our Sample Between Multiple Lines

We now consider in detail sampling from multiple lines within a consignment. Suppose that the regulator believes it to be appropriate to sample across the *K* lines of a consignment as though they were a single mixed line. While we accept that each line might have a different prevalence, our criterion is that the overall prevalence in the consignment is equal to the design prevalence.

We shall find which combination of line prevalences (that satisfy the design prevalence) corresponds to the smallest overall sensitivity. By basing our calculation of the total number *n* of samples required on that combination of prevalences, we will ensure that the sensitivity of the inspection will be always greater than the required design sensitivity, *S_d_*.

We shall sample a proportion *w_k_* of the total sample from line *k*. Hence the sample size per line is *n_k_* = *w_k_n*, such that ∑_*k*_ w_k_ = 1. There are *N_k_* units in the *k*^th^ line making a total of *N* = ∑_*k*_ N_k_

If there are *d_k_* contaminated items in line *k* we could use the Hypergeometric distribution to calculate the probability that none of these would be found. The result is mathematically intractable, and it is both more convenient and more conservative to use the Binomial approximation based on a contamination rate expressed as a proportion of 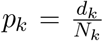. The joint contamination rate, *p* (our design prevalence), satisfies ∑_*k*_ N_k_p_k_ = *N* · *p* = ∑_*k*_ d_*k*_.

When sampling from multiple lines, the sensitivity *S* of the inspection is of the same form as Equation (1), namely

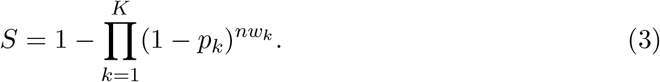

Minimising Equation (3) is equivalent to maximising ∑_*k*_nw_k_ log(1 –*p_k_*), subject to the constraint placed by the joint contamination rate, ∑_*k*_ ·N_k_p_k_ = *N p*. It is straightforward to show by the method of Lagrange Multipliers (Lagrange, 1811) that the combination of *p_k_* for which the sensitivity is least is: 
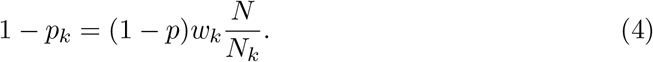

We can now consider the optimal values for the weights *w_k_*, beginning with the best choice, which is splitting the sample proportional to the line sizes.

## 2.2 Dividing the Sample Size Proportional to the Line Sizes

In Equation (4), if we choose *w_k_* = *N_k_/N* then *p_k_* = *p* minimises the sensitivity, *S*.

We choose n by substituting these values into Equation (3), giving 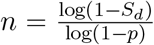. This choice of *n* and weights *w_k_* = *N_k_/N* ensure that the realised sensitivity will be no worse than the design sensitivity, irrespective of the individual line prevalences that satisfy the design prevalence.

The total sample size is the same as if we were sampling from a homogeneous population, as evidenced by the finding that having the same prevalence in each line corresponds to the combination of prevalences that gives the minimum sensitivity if we choose our weightings to be proportional to the line size. For any other combination of line prevalences that overall meet our design prevalence, the sensitivity of the inspection will be greater than the design sensitivity.

Figure 1 compares proportional and non-proportional allocation by way of an example of a consignment with two lines, one line has 20000 units, the other has 10000. We wish to find contamination present at the design prevalence of 0.5%, with 95% sensitivity. As already mentioned this requires a 600 unit sample (which actually corresponds to a 95.06% sensitivity). Consider three allocation schemes: the proportional allocation as just derived, requiring a sample of 400 units from the first line and 200 units from the second, and two non-proportional schemes where the sample sizes in each line are 395/205 units and 405/195 units respectively.

**Figure 1:**
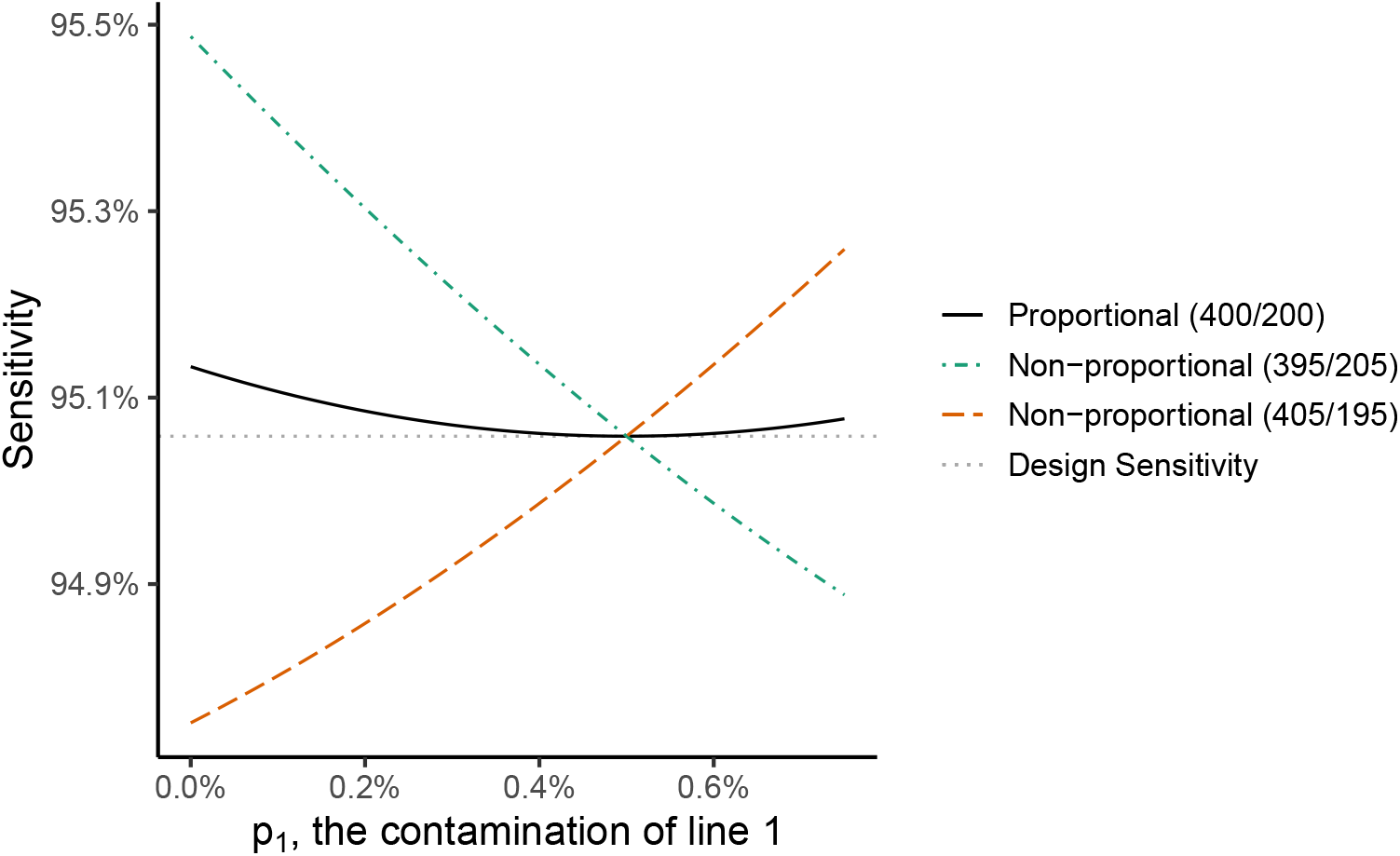
Achieved sensitivity under different sample allocation schemes. If each line has the design prevalence of 0.5%, the desired sensitivity of 95.06% (the grey horizontal line) requires a total sample size of 600 units. The figure shows the sensitivity obtained from different divisions of the 600 units between lines when the prevalence in line 1 and line 2 vary such that the overall prevalence is 0.5%. If we split the 600 samples proportionally, the solid black line shows the sensitivity obtained is always greater than the desired sensitivity. For a non-proportional allocation, the sensitivity is sometimes greater and sometimes less than desired.

Figure 1 demonstrates the achieved sensitivity that would result from each allocation scheme as a function of the true contamination rate of the first line. The solid line shows the achieved sensitivity if we used proportional allocation, the horizontal line shows the nominal sensitivity, and the other lines show the two sensitivities achieved by the non-proportional allocation schemes. The key feature to note in Figure 1 is that the achieved sensitivity is *always* greater than the nominal sensitivity of 95% under proportional allocation, whereas it may be less under non-proportional allocations for some prevalence combinations that meet the design prevalence.

## 2.3 Variations of the Problem

There are a number of minor variations to the problem of splitting the sample size between a number of lines. The derivations are not given but follow a similar method to the above.

### 2.3.1 Imperfect Inspection

Sometimes our inspection will not be fully effective, and we have a probability *e_k_* that inspection of a contaminated item in line *k* will detect the contamination. When our inspection method is less than perfect, we need to take more samples to compensate. Define *M_k_* = *N_k_/e_k_*. If we assume that our sample is divided between lines in proportion to *M_k_*, we can show that the minimum sensitivity occurs when the *apparent* prevalence (*p_k_e_k_*) in each line is the same by using the method in Section 2.1. From that we find that the number of samples required should be based on an adjusted (smaller) prevalence 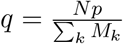 to give 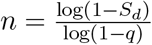.

### 2.3.2 Design Prevalence as an Absolute Number

Occasionally the design prevalence is specified as an absolute number *D* of contaminated items. Replacing *p* by 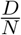 in the above gives the required sample size which, as before, would be split proportionally between the lines: 
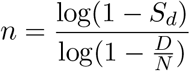

For an absolute design prevalence, log(1 *− D/N*) needs to be calculated for each consignment. To simplify this, one can increase the sample size slightly by using the approximation ln (1 *− D/N*) *≈ −D/N* (which is equivalent to using the Poisson approximation to the Binomial). This gives the overall number sampled being proportional to the number in the consignment: 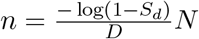.

### 2.3.3 Not Knowing Line Sizes Accurately

So far we have assumed that the counts for each line are accurately known. If the percentage errors in the counts are likely to be similar, this will be of little concern, since the relative contribution each line makes to the total will stay much the same. If, however, there is more uncertainty, the number of samples required needs to be increased for each line.

Suppose that we think the actual line sizes could be between *N_k_*(1*−α_k_*) and *N_k_*(1+*β_k_*). The consignment size would be between *N* (1*−α*) and *N* (1+*β*), the sum of the lower and upper line sizes respectively. Hence the weighting for line *k* should lie between 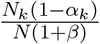 and 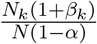. To be conservative, we use the upper limit of this range to determine the number of samples per line in terms of *n* calculated based on Equation (2) using our desired sensitivity and design prevalence: 
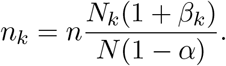

Our uncertainty about line size means that we need to take more samples in total, 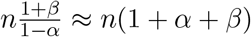. As an example, if our uncertainty of the size of the consignment was of the order of *±*10%, then we need to increase the sample size by approximately 20%.

### 2.3.4 Using Fixed Sample Sizes

Regulators might wish to choose fixed sample sizes for each line, rather than allocate sample sizes proportional to the line sizes. For example, we could take an equal number of samples from each line. However, for such weightings, more samples are required in order to ensure the design sensitivity *S_d_* is met. For all practical purposes, the number of samples (*m*) required for fixed sample sizes has to be chosen so that for each line the number of samples taken, say *m_k_* = *w_k_m*, is greater than or equal to 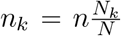, the number of samples required if proportional weightings had been used.

## 3 Discussion and Conclusions

A truly random sample from the entire consignment will give the desired sensitivity regardless of any clustering of contamination in the consignment. This would result in a random number of samples being taken from each line—possibly only a few or even none—something regulators might be uncomfortable with.

We have shown how a standard sample size may be split between a mixed-line consignment, while still giving the desired chance of detecting contamination if it is present at a specified rate for the entire consignment. The critical point for exporters to understand is that if contamination is found in just one line, the entire consignment has not satisfied the import requirements and would be deemed to have failed the inspection with the resultant consequences.

The reverse is true for regulators: it is important that they do not deem only the lines in which contamination was found as non-compliant and accept the rest. The lines in which no contamination has been found have not had sufficient inspection to demonstrate that they meet the design sensitivity and prevalence requirements. While it is out of scope for this paper, it must be pointed out that simply taking more samples to make up the difference in the ‘clean’ lines to the total sample size (e.g. 600 units) is not enough. In designing the total sample size required, the fact that we will tolerate finding a contaminated unit must be taken into account.

We note that there are reasons for which treating lines separately makes operational sense. One is that the products may carry different kinds of pests that themselves present different risks. Another is that the exporter may not wish to take the chance that contamination in one line will affect the treatment of all of the lines in the consignment. Some other practical considerations need to be made, but these can usually be resolved by taking additional samples either in some lines only or by increasing the number of samples required in total.

Our result relies on the assumption of exact proportional allocation of the samples to lines based on their counts. In some situations, the number of units in a line might differ from the nominal count, so that an exact proportional allocation would not be made. We have shown that increasing the sample size in proportion to the likely variation provides a way to ensure that the desired sensitivity is still met.

Furthermore, our result assumes that the sampling is done randomly within each line. If contamination is likely to be clustered and the sampling is not random (for example inspecting all fruit within a number of randomly-selected boxes) a different method must be used to determine the sample size (e.g. Venette et al., 2002). Extending such results from a single line is outside the scope of this paper.

Using a proportional allocation of the sample might not be prudent when the number of items in one line greatly exceeds the number in the other lines. An example of this might be with one line being melons, and one of the other lines being cherries. The problem is that proportional allocation might result in only one or two units being selected from lines with few units. While the lines with few units might only contribute a small proportion of the contamination, there may be misgivings that they haven’t been adequately inspected. One way this could be resolved is by considering them to be, from the point of view of sampling, as two separate consignments. Another alternative might be to consider a box of cherries as the unit, which might give comparable unit numbers in the lines.

Another solution might be to top up the calculated number of samples to make a minimum sample per line. This would guard against missing gross contamination in a line with few units which, while not contributing greatly to the overall contamination, would be of concern if present. For example a minimum sample of 30 in a line would detect a contamination rate of 10% in that line with a 95% probability. The other advantage in having a minimum sample size would be that information about that particular item type or source would be more quickly accumulated.

If the types of contamination in some lines are thought to have greater consequences than others, one could take extra samples above what is required in those lines, for example take twice as many. While taking extra samples is a form of non-proportional allocation, it is based on the number determined by proportional allocation: taking extra samples above the proportional allocation would increase the sensitivity of the inspection. However, to ensure the design sensitivity is met for a more general division of the sample numbers between lines (such as equally between the lines), no line should have fewer samples taken from it than the number determined by proportional allocation.

Finally, it cannot be emphasised enough: when the sample is stratified proportional to the stratum size, if contamination is found, even if it is in just one line, the whole consignment has to be deemed non-compliant and subject to whatever requirement non-compliance imposes. If this is not acceptable, then individual lines (or groups of lines) must be inspected separately, with each component subject to the specified compliance test.

## References

Cochran, W G (1977). Sampling techniques. 3rd. John Wiley & Sons, Inc.

Colautti, Robert I, Sarah A Bailey, Colin D A van Overdijk, Keri Amundsen, and Hugh J MacIsaac (2006). “Characterised and Projected Costs of Nonindigenous Species in Canada”. In: Biological invasions 8.1, pp. 45–59. issn: 1387-3547, 1573-1464. doi: 10.1007/s10530-005-0236-y.

Giera, N and B Bell (2009). Economic Costs of Pests to New Zealand. Tech. rep. 2009/31. Biosecurity New Zealand, Ministry of Agriculture and Forestry.

Hoffmann, Benjamin D and Linda M Broadhurst (2016). “The economic cost of managing invasive species in Australia”. In: Working paper series 31, pp. 1–18. issn: 0898-2937. doi: 10.3897/neobiota.31.6960.

International Plant Protection Convention (2008). International standards for phytosanitary measures: ISPM 31, methodologies for sampling of consignments. Food and Agriculture Organization of the United Nations, Rome, Italy.

Lagrange Joseph-Louis (1811). Mécanique Analytique. Paris, Ve Courcier.

Pimentel David (2011). Biological Invasions: Economic and Environmental Costs of Alien Plant, Animal, and Microbe Species. Second. Hoboken: CRC Press, 2011.

Robinson, Andrew P (2017). Compliance and risk-based sampling for horticulture exports. Tech. rep. 1501E, Deliverable 7. Centre of Excellence for Biosecurity Risk Analysis.

Venette, Robert C, Roger, D Moon, and William, D Hutchison (2002). “Strategies and statistics of sampling for rare individuals”. en. In: Annual review of entomology 47, pp. 143–174. issn: 0066-4170. doi: 10.1146/annurev.ento.47.091201.145147.

